# Environmental factors that impact the development of infective juveniles of entomopathogenic nematode *Steinernema hermaphroditum*

**DOI:** 10.64898/2026.04.07.717109

**Authors:** Mengyi Cao

## Abstract

Animals sense and integrate complex external cues to make developmental decisions that help them better survive and adapt to their natural habitats. Under environmental adversity, nematodes can enter an alternative developmental pathway to form a diapautic and stress-resistant stage, termed the dauer larvae. While dauer formation has been well characterized in *Caenorhabditis elegans*, how environmental factors influence the development of analogous stages in parasitic nematode species remains largely unexplored. This study examines how symbiotic bacteria, temperature, and pheromones affect the formation of the infective juvenile (IJ), a dauer-like and infective stage, in the insect-parasitic nematode *Steinernema hermaphroditum*. In contrast to *C. elegans,* where dauer entry requires exogeneous pheromone and is promoted by heat, IJ development in *S. hermaphroditum* does not rely on exogeneous pheromone and is enhanced by reduced temperature. Moreover, the presence and absence of live symbiotic bacterium *Xenorhabdus griffiniae* functions as an ON-and-OFF switch that regulates IJ formation. Nutrient-rich liver-kidney media that mimics host insect environment showed IJ entry induction in a DAF-22 dependent manner, suggesting that short-chain ascarosides (pheromone) promote *S. hermaphroditum* IJ formation through a conserved pathways across nematode species. Together, these findings reveal that environmental cues, such as temperature, microbial diet, and pheromone, can be perceived differently or similarly by *S. hermaphroditum* compared with *C. elegans*, reflecting species-specific adaptations to distinct ecological niches and life history strategies.

**Summary Statement:** Parasitic nematodes rely on infective stages for host invasion, yet their developmental triggers remain largely unexplored. This study reveals how environmental cues regulate IJ formation.

## Introduction

Animals enter diapause, a type of dormancy state, to adapt to harsh environmental conditions such as droughts, winter, and starvation. By sensing and integrating external signals, the animals can respond with an endogenous regulatory program that leads to developmental decisions to enter diapautic arrest (Hand et al. 2016). Under favorable conditions, free-living nematodes proceed through a reproductive cycle, growing and molting through four larval stages before becoming adults that lay eggs (Fig. 1C). However, when conditions become unfavorable, they can switch to an alternative developmental pathway that leads to a diapause stage known as the dauer larva. The dauer larvae exhibit distinct morphological and physiological traits compared to reproductive stages: they are non-feeding due to closed mouth and anus, and a collapsed intestine (Fig. 1D). They are developmentally arrested and highly resistant to environmental stress. Dauer entry is triggered by adverse cues such as non-optimal temperatures, overcrowding, or a combination of dauer pheromone accumulation and starvation, indicated by the absence of food signals (Golden and Riddle 1984). Since its discovery, the free-living model nematode *Caenorhabditis elegans,* naturally inhabited in rotten fruits, has been extensively studied for its dauer biology, offering critical insights into the molecular mechanisms that govern developmental transitions into diapause (Fig. 1C and 1D) (Hu 2007).

**Figure 1:**
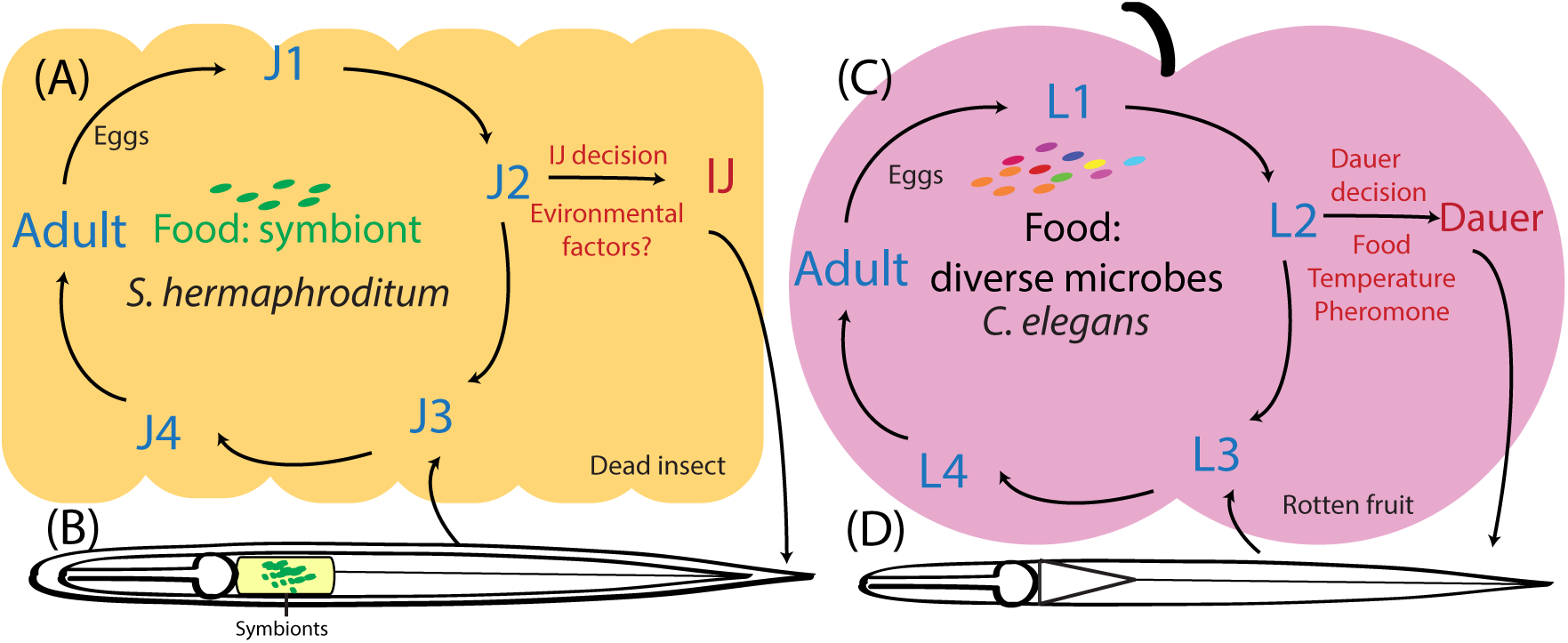
Life cycle and diapause decision making in two nematode species. (A): Insect-parasitic nematode *S. hermaphroditum* feeding on symbiotic bacterium *X. griffiniae* in the insect cadaver and develop through reproductive (blue) or diapautic/IJ (red) pathways. (B): An infective juvenile (IJ) stage of nematode carrying symbiotic bacteria in the intestinal pocket. (C)*: Caenorhabditis elegans* reproduction (blue) and dauer developmental pathway (red) on a rotten fruit. (D): A dauer *C. elegans* nematode at dispersal stage.

Research on *C. elegans* dauer signaling pathways is highly reliant on laboratory experiments and has provided important insights into the molecular mechanisms underlying similar physiological states in other nematodes, including those involved in cryobiology and parasitism. For example, dauer has served as a comparative model to study cryptobiosis in an ancient *Panagrolaimus* nematode reanimated after being frozen in Siberian permafrost for approximately 46,000 years (Shatilovich et al. 2023). One theory, known as the “dauer hypothesis,” proposes that the dauer stage is a prerequisite for the evolution of parasitism (Crook 2013; Vlaar et al. 2021). A logical extension of this hypothesis is that conserved molecular pathways underlie dauer development in both free-living and parasitic nematodes.

Despite the appeal of this model and evidence supporting conserved roles for certain signaling pathways, *C. elegans* dauer biology does not fully capture the unique ecological and developmental complexities of diapause regulation in parasitic species. Free-living and parasitic nematodes likely possess distinct sensory modalities and molecular programs governing dauer entry, which limits the extent to which *C. elegans* can serve as a universal model. Nevertheless, little is currently known about the mechanisms driving dauer entry in parasitic nematodes. Although dauer-like stages have been observed in both plant- and animal-parasitic nematodes, standardized protocols remain lacking for characterizing environmental factors that influence dauer decisions in these species. Evidence for dauer in parasitic nematodes are mostly based on genomic and transcriptomic data, without *in vivo* validation (Vlaar et al. 2021).

The soil-dwelling entomopathogenic (insect-parasitic) nematodes of the genus of *Steinernema* spp. live a unique lifestyle that involves a mutualistic relationship with symbiotic bacteria. During the dauer-like stage, termed the infective juvenile (IJ), *Steinernema* nematodes carry species-specific *Xenorhabdus* symbionts within an intestinal pocket called the receptacle (Fig. 1A and 1B). IJs exhibit a repertoire of behaviors including dispersal, chemotaxis, nictation, and jumping to actively locate and invade insect hosts in the soil. Upon host entry, they release their symbiotic bacteria, which assist in killing the insect. The nematodes then reproduce inside the cadaver, feeding on their symbionts (Mucci et al. 2022). The decision to enter the IJ stage is made within the nutrient-depleting environment of an insect cadaver, where nematodes reach high population densities. While it is reasonable to hypothesize that *Steinernema* perceives and responds to environmental cues similarly to *C. elegans* in regulating dauer/IJ entry, this hypothesis has not been rigorously tested.

To investigate the development and behavior of dauer or infective juvenile (IJ) larvae, it is essential to establish controlled experimental conditions in the laboratory. In *C. elegans*, crude pheromone cocktails, termed “daumone” (dauer pheromone) extracted from liquid culture is typically used to induce dauer entry on agar plates with heat-killed *E. coli* bacteria as a controlled food source (Golden and Riddle 1982; Schroeder and Flatt 2014). Under these conditions, eggs laid by adult hermaphrodites were tested their ability to develop into either reproductive stage (J4) or dauer within 48 hours of the assay. However, comparable experimental protocols have not been established for studying IJ formation in parasitic nematodes. By adapting *C. elegans* dauer assays to *Steinernema hermaphroditum*, I examined how *S. hermaphroditum* responds to environmental cues, revealing notable differences. Unlike *C. elegans*, where elevated temperature promotes dauer entry, cold exposure enhances IJ formation in *S. hermaphroditum*, indicating opposite thermosensory responses. Other cues, such as bacterial food availability and pheromones, are also interpreted in a different manner during their respective diapause transitions. These findings contribute to the experimental evidence to examine the “dauer hypothesis,” which posits that dauer and IJ formation rely on conserved signaling pathways. The observed divergence in environmental cue perception suggests that nematodes employ distinct strategies for regulating diapause, shaped by their species-specific evolutionary ecologies. As an emerging genetic model, *S. hermaphroditum* holds great promise for uncovering novel signal transduction pathways and neural circuits underlying IJ development, thereby complementing and expanding our understanding of diapause regulation across nematode species.

## Materials and Methods

### Nematode liquid culture and crude pheromone extraction

#### *C. elegans* liquid culture

*C. elegans* liquid culture was adapted from (Schroeder and Flatt 2014). Briefly, wild-type *C. elegans* (N2 strain) was grown on Nematode Growth Media (NGM) (Stiernagle 2006). *E. coli* OP50 was grown overnight at 37°C with aeration. Each NGM plate was seeded with 50 μL of *E. coli* OP50 overnight culture and start with 1-3 gravid *C. elegans* adult hermaphrodite until the food is almost depleted at room temperature. Fifteen to twenty petri-dishes of nematodes were washed from the plates into approximately 1250 mL of S media supplemented with 0.25% supplemented with *E. coli* OP50 (pellets from 2 L *E. coli* are used for 1250 mL liquid culture). The liquid culture was shaking on an orbital shaker at 20°C, 200 rpm. Once the bacterial food was depleted in the liquid culture (4-5 days), a new batch of *E. coli* OP50 was pelleted from 2 L of overnight culture and added to feed the nematodes. *C. elegans* was grown in liquid culture to a density of 30-50 dauers per μL and used for subsequent pheromone extraction.

#### *S. hermaphroditum* liquid culture

*S. hermaphroditum* liquid culture was adapted from *C. elegans* liquid culture described above. Instead of growing nematodes in S media, LB supplemented with 5 μg/mL of cholesterol was used. Lipid agar plate (per liter: 15 g bacto agar, 8 g nutrient broth, 5 g yeast extract, 2 g MgCl_2_· 6H2O, 1 g sodium pyruvate, 7 ml corn syrup, and 4 ml corn oil (Vivas and Goodrich-Blair 2001)) were seeded with 600 μL of *X. griffiniae* culture which grew into bacterial lawns at room temperature. *S. hermaphroditum* nematodes were grown on bacterial lawns until almost starve. Approximately 10-15 plates of *S. hermaphroditum* on Lipid Agar plates were used to start a total 1250 mL of liquid culture. Liquid culture was supplemented with 30 μg/mL Chloramphenicol to inhibit bacterial overgrowth and added *X. griffiniae* pellet from 2 L culture and shaking at 25.5 °C, 200 rpm. Once the bacterial food was depleted in the liquid culture on day 6 clearance of turbidity, 2 L of *X. griffiniae* was spun down and the pellet was added to the liquid culture to feed the nematodes. The feeding was performed twice. IJ density reached 15-20 IJs per μL (15,000 – 20,000 IJs per mL) when the liquid culture was used for pheromone extraction.

#### Pheromone extraction from liquid cultures

the method to extract crude pheromones from liquid cultures was adapted from (Schroeder and Flatt 2014). Briefly, liquid cultures were used at a density of approximately 30-50 dauers per μL (30,000 – 50,000 dauers per mL) for *C. elegans*, or 10 IJs per μL (10,000 IJs per mL) for *S. hermaphroditum*, with a total 1250 mL of liquid culture for each species. The supernatants of liquid cultures were acquired by centrifugation, filtered through 0.4 μm microporous membranes, boiled to remove water, and eventually crusted in a 50 mL glass beaker. Pheromone was extracted by pouring 100% ethanol over the crust. Once the color of ethanol extract changed to yellow or brown, indicating the presence of pheromone in the extract, the supernatant was collected in 50 mL conical tubes stored at −20°C. The ethanol extraction was repeated with the same crust for multiple times over 2-4 days. The ethanol extracts were combined and heated in a glass beaker under 50°C with stirring using a heat plate under the fume hood. Pheromone extract was resuspended in sterile water, filtered-sterilized by passing through 0.2 μm microporous membranes, and kept at −20°C.

### Dauer and IJ entry assays

#### Preparation of live and killed bacteria

*E. coli* OP50 liquid culture in was grown in LB media at 37°C overnight with aeration. Approximately 20 mL overnight culture was pelleted by centrifugation, washed twice with S Basal media (Stiernagle 2006), and adjusted to 8% (w/v) in S Basal. The 8% bacterial resuspension were aliquoted in microfuge tubes and heat-killed at 95-100°C heat blocks for 5 minutes. The 100% killing efficiency was confirmed by streaking 2 μL of killed bacteria onto LB agar plates and incubate at 37°C overnight.

For *S. hermaphroditum* IJ entry assays, *X. griffiniae* (HGB2511) overnight culture was grown at 30°C, spun down by centrifugation, washed in S-basal for twice, and adjusted to 4% or 8% (w/v). Live bacteria were supplemented with 30 μg/mL Chloramphenicol and used immediately for IJ entry assays on NG media (per 250 mL per bottle: 0.3 g NaCl, 2.5 g Bacto-agar, 97.5 mL distilled water, autoclave, supplemented with 100 μL 5 mg/mL cholesterol, 100 μL 1M CaCl_2_, 100 μL 1M MgSO_4_, and 2.5 mL 1M KPO_4_ PH 6). The NG media was also supplemented with 30 μg/mL Chloramphenicol to c growth. Heat-killed bacteria were prepared the same as heat-killed *E. coli* OP50 as described above. Cold-killed bacteria were stored at 4°C for one week, confirmed killing efficiency by streaking out bacteria and grew at 30°C, then used in IJ entry assay.

#### *C. elegans* dauer entry assay

*C. elegans* was grown on NGM agar plates seeded with *E. coli* OP50 without starving. Pheromone plates were freshly prepared the night before experiment by melting and transferring 2 mL of NG agar (supplemented with 5, 10, 15, 20 μL/mL pheromone extracted from *C. elegans* N2 strain), dried overnight, and used immediately the next day. On the day of the experiment, 5 μL of live *E. coli* OP50 overnight culture was seeded at the center of the pheromone plate. Ten well-fed adult *C. elegans* were placed on each pheromone plates and allowed to lay approximately 75-100 eggs, then the adults were removed from the pheromone plates, and 20 μL of heat-killed *E. coli* (8% w/v) was added to each pheromone plate until dried. The pheromone plates were sealed with parafilm, incubated at 25.5°C for 24-48 hours and monitored every 12 hours. Once dauer decision is made, the number of dauer and non-dauers (L3, L4, adults) were scored based on morphology. Each agar plate is considered one biological replicate. For each replicate, approximately 50-100 nematodes were scored, and the percentage of dauer entry was calculated based on the number of animals that entered the dauer stage. The same batch of pheromone was stored at −20°C and used in all independent experiments.

#### *S. hermaphroditum* IJ entry assay with exogeneous pheromone

*S. hermaphroditum* were propagated on NGM agar plates seeded with sufficient amount of symbiotic bacteria, *X. griffiniae,* to keep them from starving. The night before experiment, IJ entry assay plates freshly prepared by melting and transferring 2 mL of NG agar supplemented with 30 μg/mL Chloramphenicol and crude pheromone extracted from wild-type *S. hermaphroditum* (0 μL, 15 μL, 30 μL or 45 μL per 2 mL). On the day of the experiment, 10-20 gravid wild-type hermaphrodites were placed onto each agar plate seeded with 5 μL fresh overnight culture of symbiotic bacteria until 50-100 eggs were laid. 20 μL of 4% or 8% (w/v) symbiont was prepared from freshly grown overnight culture (live bacteria), heat-killed, or cold-killed (refrigerated) bacteria (described above in the ‘preparation of live and killed bacteria’ section), then added to the embryos on each agar plate. The assays were incubated at the respective temperatures (20-30°C) until the IJ decision was made (2-4 days). The IJs and non-IJs were scored based on morphology. For each replicate, approximately 50-100 nematodes were scored, and the percentage of dauer entry was calculated based on the number of animals that entered the IJ stage. The same batch of pheromone was stored at −20°C and used in all independent experiments.

#### *S. hermaphroditum* IJ entry assay on Liver-Kidney agar plates

to perform IJ entry assay on liver-kidney agar pates, approximately 200 axenic embryos were extracted and grown on liver-kidney agar seed with *X. griffiniae* lawn and incubated at 25.5°C (Vivas and Goodrich-Blair 2001; Murfin et al. 2012). On days 8-10 of incubation, the second-generation progeny are forming IJs that migrate to the edge of the plate. Approximately 100 - 200 μL of sterilized water was added to the inner rim of each Liver-Kidney agar plate and agitate thoroughly to resuspend the nematode population close to the edge and on the rim of the plate where IJs are most concentrated and ready for migration. Nematode population collected from each plate was washed and incubated in sterilized water overnight. IJs survive in water while non-IJs are killed, leaving visible nematode carcass. Approximately 100 – 200 nematodes (including carcass) per sample was examined under dissecting scope to determine if they are IJ (live) or non-IJs (dead) by morphology and viability. Percent IJs were calculated using number of IJ out of total number of nematodes. The *S. hermaphroditum* nematode strains used in this experiment was wild-type (PS1961(Cao et al. 2021)), *Sh-daf-22* mutant (MC0006 (Cao 2023)), and *Sh-unc-22* mutant (MC0003 (Cao 2023)).

#### Statistical analysis

Statistical analysis (Student’s *t-test*, One-way and Two-way ANOVA tests) were performed using GraphPad Prism 11. Significant differences were shown as \**P<0.05, **P<0.01, ***P<0.001, and ****P<0.0001*.

## Results

### Symbiotic bacteria regulate IJ development in the absence of exogeneous pheromone

In *C. elegans* dauer decision assays, a gradient of exogeneous nematode pheromone is required to induce dauer formation in a dose-dependent manner (Fig. S1A). Under optimized conditions in a dauer entry assay, the synergistic effects from the presence of pheromone and limited food source, usually heat-killed *E. coli* OP50, can induce approximately 50% of wild-type embryos enter the dauer stage and 50% proceed to the reproductive (L4) stage, mimicking a binary developmental decision (Fig. S1A). In the absence of pheromone, embryos grown on heat-killed *E. coli* exhibit 0% dauer entry (Fig. S1A).

To adapt *C. elegans* dauer entry protocol for the *S. hermaphroditum* infective juvenile entry assay (Fig. 1A), I first compared live and heat-killed symbiotic bacteria (*Xenorhabdus griffiniae*) to identify conditions under which approximately 50% of embryos develop into IJs (Fig. S2A). In contrast to *C. elegans*, S. *hermaphroditum* embryos can develop into IJs in vitro without exogeneous pheromone (Fig. 1 and Fig. S2). Approximately 50% IJ entry was achieved using 5 µL of live *X. griffiniae* without exogeneous pheromone, whereas increasing the amount of live symbiotic bacteria progressively suppressed IJ formation in a dose-dependent manner (Fig. S2A). IJ formation was 27.9% and 2% on 7.5 µL and 15 µL of 4% live symbiotic bacteria, respectively (Fig. 1B and 1C). In contrast, IJ formation increased to 90.2% and 79.3% heat-killed symbiotic bacteria, respectively (Fig. 1B and 1C). Together, these findings demonstrate that the absence of live symbiotic bacteria functions as a binary ON/OFF switch that regulates IJ development in *S. hermaphroditum* (Fig. 2D). Cold-killed (refrigerated) *X. griffiniae* also induced IJ formation (Fig. S2B), although the response was more variable than that observed with heat-killed counterparts (Fig. S2C). This finding suggests that live symbiotic bacteria may produce heat-sensitive and volatile cues that promotes *S. hermaphroditum* reproduction and suppressing IJ formation.

**Figure 2:**
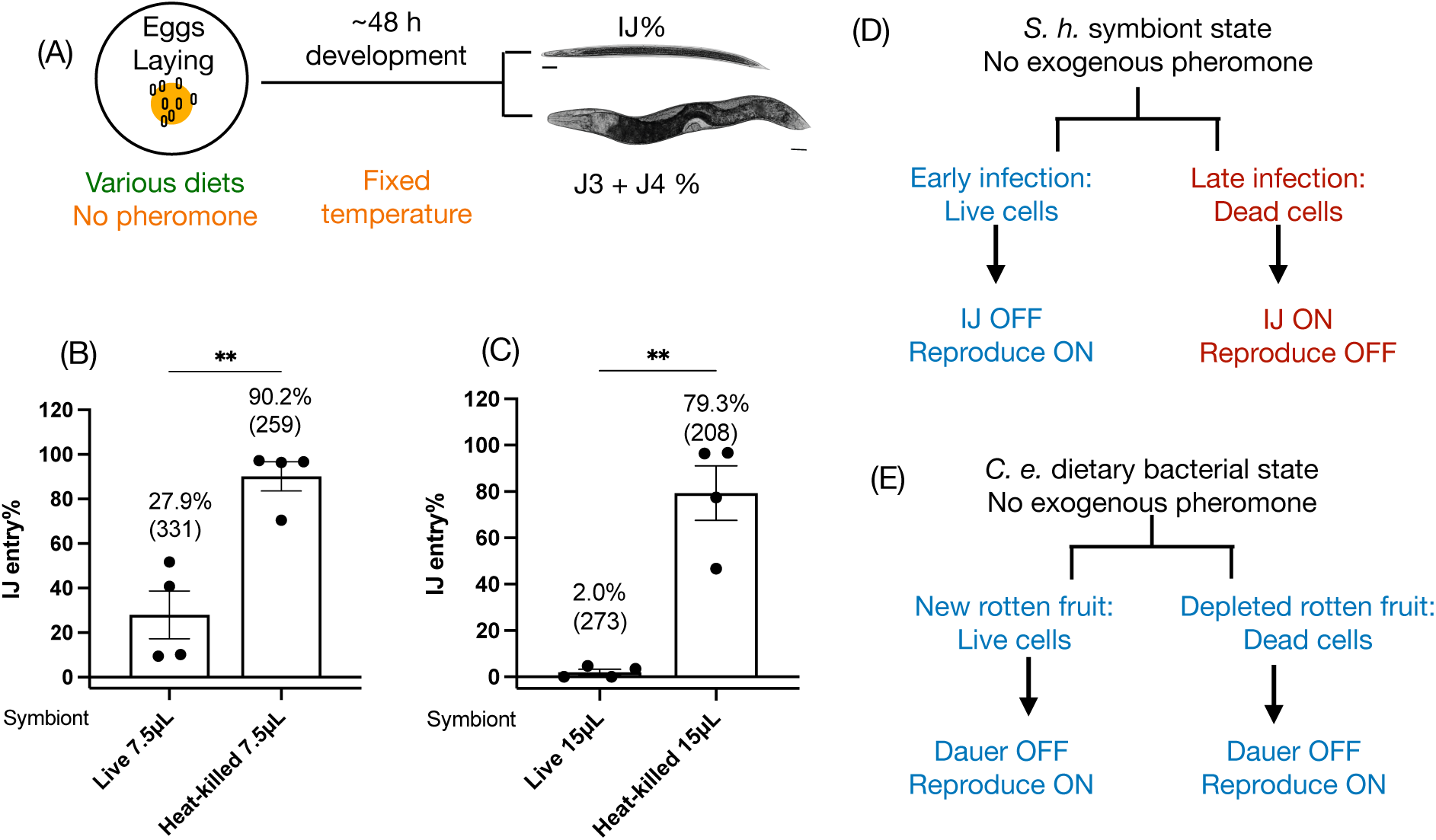
Effects of live and heat-killed symbiotic bacteria on *S. hermaphroditum* IJ entry. (A): A schematic diagram of IJ entry experiments. Adult nematodes were transferred onto NG agar. Approximately 50-100 *S. hermaphroditum* eggs were freshly laid for each plate incubated at the 25.5°C until the IJ decision was made (24-48 hours). (B and C): IJ entry on NG agar supplemented with 30 μg/mL Chloramphenicol and 7.5 μL (B) or 15 μL (C) of live or heat-killed symbiotic bacteria *X. griffiniae* (8% w/v). Each data point represents a biological replicate from two independent experiments. Statistical analysis was performed using t test. The average percent IJ entry is shown. Number in parenthesis shows total number of animals (sample size) in each condition. (D and E): Hypothetical models explaining how symbiotic bacteria signal *S. hermaphroditum* IJ developmental decision (D) in comparison to *C. elegans* dietary *E. coli* signaling dauer decision.

**Figure 3:**
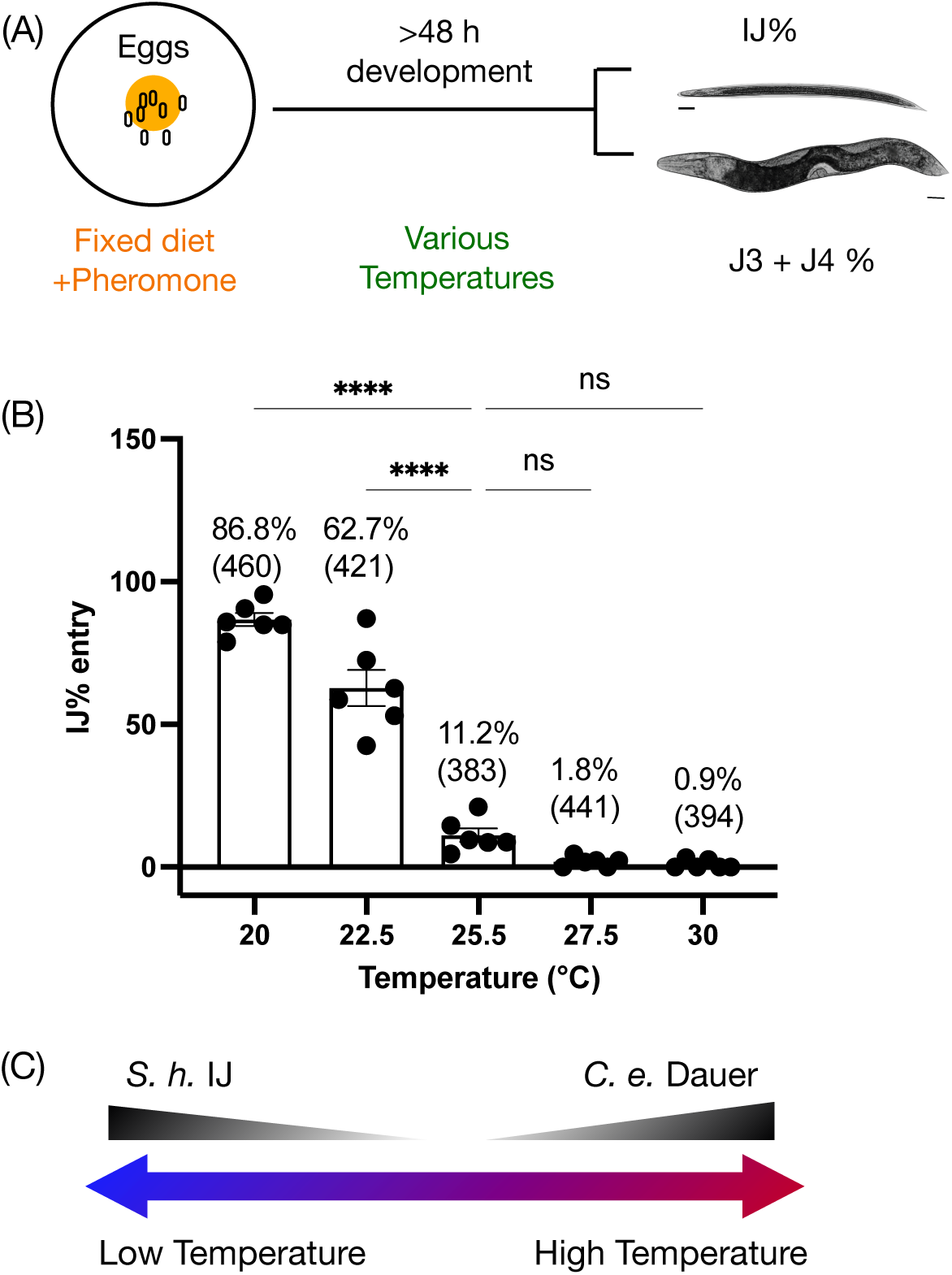
The impact of temperature on *S. hermaphroditum* IJ entry. (A): A schematic diagram: the IJ entry assay was performed with supplementation of crude pheromone (15 μL per mL agar). Each pheromone plate contains 15 μL of freshly prepared live symbiotic bacteria *X. griffiniae (*4% w/v) and treated with 30 μg/mL Chloramphenicol, then added to 50-100 embryos on each agar plate. The assays were incubated at the respective temperatures (20-30°C) until the IJ decision was made in 2-4 days. (B): Each data point represents an individual plate. Three independent experiments were performed. Statistical analysis was performed using One-way ANOVA. The average percentages of IJ entry and standard errors are shown. Number in parenthesis indicate total number of animals (sample size) in each condition. (C): A schematic diagram summarizing the impact of temperatures on the development of *S. hermaphroditum* IJ and *C. elegans* dauer.

Overall, these results are consistent with the natural progression of insect infection, during which *Xenorhabdus* symbionts initially proliferate to form a nearly monoxenic bacterial population. During this early phase, when live symbiotic bacteria are abundant, nematodes primarily undergo reproductive development. As infection progresses and dead bacterial cells accumulate, nematodes are induced to enter the infective juvenile (IJ) stage for dispersal (model presented in Fig. 2D). In contrast, free-living *C. elegans*, which inhabits environments containing diverse microbial communities, is observed to rely primarily on species-specific pheromones to coordinate dauer entry rather than on the physiological state of a particular bacterial food source (Fig. 2E).

### Temperature regulates *S. hermaphroditum* IJ formation and *C. elegans* dauer formation in the opposite directions

Next, I tested whether temperature influences *S. hermaphroditum* IJ development. In *C. elegans*, elevated temperatures promote dauer formation (Ailion and Thomas 2000), and I initially hypothesized the same would hold true for *S. hermaphroditum*. The optimal growth temperature for *S. hermaphroditum* is at approximately 25°C (Cao et al. 2021). I performed IJ entry assays at five temperatures of 20°C, 22.5°C, 25°C, 27.5°C, and 30 °C (Fig. 2). In contrast to *C. elegans*, IJ entry in *S. hermaphroditum* was **negatively** regulated by temperature. Cold exposure at 20 °C and 22.5 °C induced approximately 86.8% and 62.7% IJ entry, respectively, on plates containing live symbiotic bacteria and crude pheromone (Fig. 2B). IJ formation progressively decreased as the temperature increased. At 25.5°C, IJ entry significantly decreased to 11.2%. As temperatures increase to 27.5°C and 30°C, percentage of IJ formation was 1.8% and 0.9%. These results demonstrate that, unlike *C. elegans*, cold exposure rather than heat, promotes IJ development in *S. hermaphroditum*.

### Low temperature and exogeneous pheromone synergistically induce IJ formation

To determine whether temperature and pheromone act synergistically to regulate *Steinernema* IJ development, I performed IJ entry assays at five temperatures using four concentrations of crude pheromone at each temperature (Fig. S3). At lower temperatures (20 °C and 22.5 °C), the presence of pheromone significantly enhanced IJ formation by approximately 33% and 26%, respectively, compared with the no-pheromone control. In contrast, at higher temperatures (25.5 °C, 27.5 °C, and 30 °C), exogenous pheromone increased IJ entry by only 11.2%, 1.8%, and 0.9%, respectively, and these increases were not significantly different from the no-pheromone control. Together, these results indicate that crude *S. hermaphroditum* pheromone and low temperature act synergistically to promote IJ formation (Fig. S3). Notably, unlikely *C. elegans* (Fig. S1A), IJ entry did not exhibit a clear dose-dependent response across the range of pheromone concentrations tested (Figs. S1B and S3).

### DAF-22 regulates IJ formation on nutrient-rich liver-kidney assays

It is possible that pheromone extraction from *S. hermaphroditum* liquid culture and IJ entry assay on NGM (nutrient-limiting) agar plate do not be the most physiologically relevant condition for evaluating the role of pheromone in IJ development. To address this possibility, I established an IJ developmental assay using liver-kidney agar plates (Fig. 4). The liver-kidney agar was prepared from grounded beef liver and kidney, a nutritious media that supports *Steinernema* reproduction mimicking their natural life cycle in an insect cadaver (Fig. 1A (Cao et al. 2021)). *S. hermaphroditum* nematodes were cultured on abundant symbiotic bacteria at permissive reproductive temperature of 25.5°C for approximately two generation (7-10 days), during the second-generation progeny development, the high nematode density and pheromone accumulation would naturally induce IJ formation (Fig. 4A). To determine whether IJ formation was specifically induced by pheromone rather than by other environmental factors, such as depletion of live symbiotic bacteria, I used a *S. hermaphroditum daf-22* mutant as control (Fig. 4B and 4C). DAF-22 is a highly conserved enzyme in the nematode pheromone biosynthesis pathway that catalyzes the final step of peroxisomal β-oxidation required for short-chain ascaroside production (Fig. 4C). Consistent with a major role for pheromone biosynthesis, the *daf-22* null mutant, which is expected to be severely deficient in ascaroside production, exhibited defects in IJ entry on liver-kidney agar compared to wild-type (Fig. 4B and 4C). In contrast, the *unc-22* mutant, used as a co-CRISPR marker control, displayed wild-type level of IJ development (Figs. 4B and 4C). By day 10, both wild-type and *unc-22* animals reached approximately 80% IJ development, while *daf-22* mutants showed approximately 30% IJ development (Fig. 4C). These results demonstrate that DAF-22-dependent pheromone biosynthesis is required for normal IJ development in *S. hermaphroditum*. the pheromone-dependent defect in IJ development observed on liver-kidney agar at 25.5 °C (approximately 60% reduction) was substantially greater than that observed on nutrient-limiting NGM agar under the same temperature conditions (<12% reduction; Figs. S3). Together, these findings demonstrate that pheromone produced under nutrient-rich, insect cadaver-like conditions is sufficient to drive robust IJ formation even at a temperature that is otherwise permissive for reproduction.

**Figure 4:**
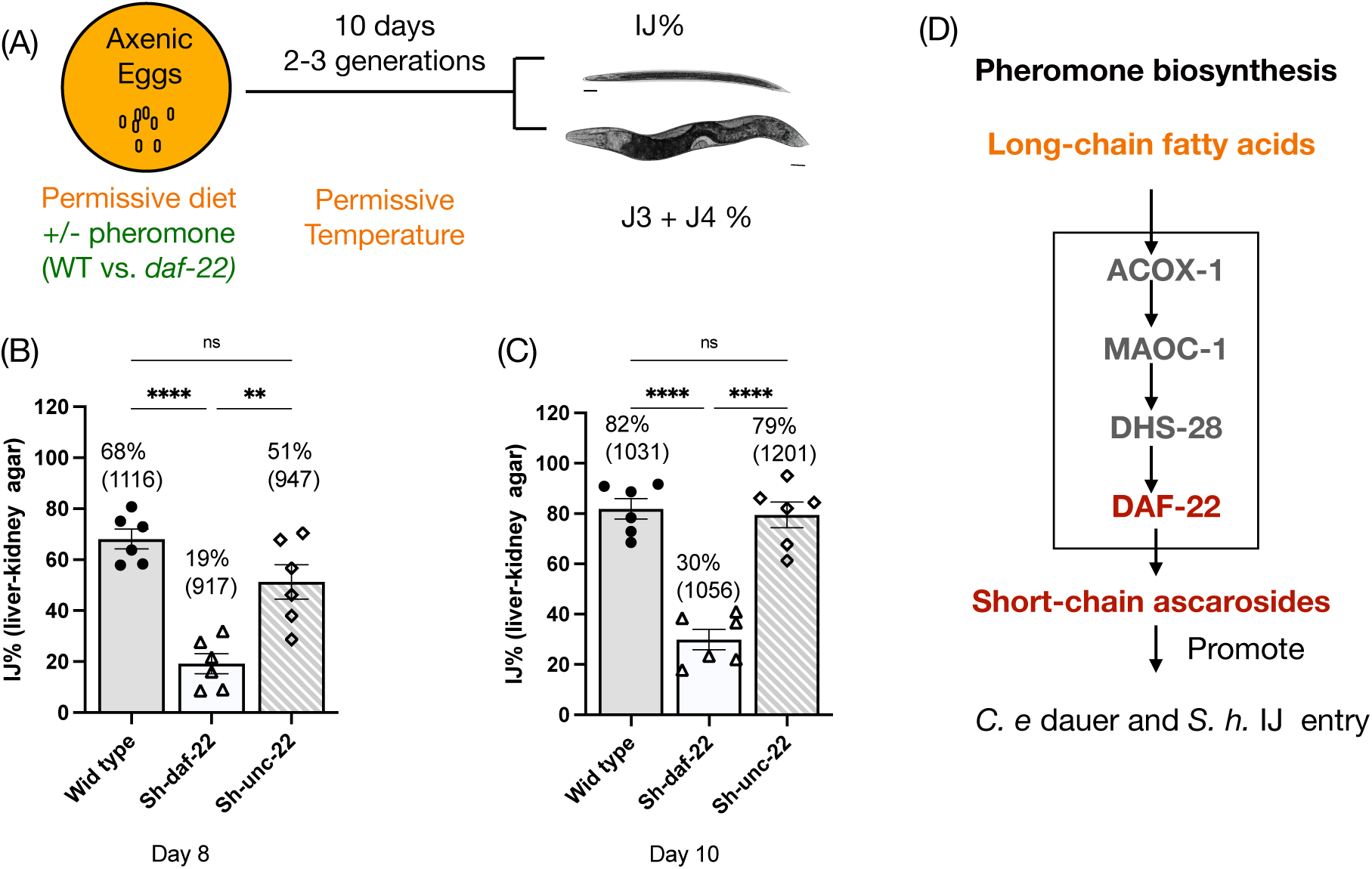
*S. hermaphroditum* IJ development on Liver-Kidney agar plates. (A): A schematic diagram: each Liver-Kidney agar plate was seeded with 600 μL live symbiotic bacteria *X. griffiniae* and grew into a confluent bacterial lawn. Axenic eggs of wild-type, *daf-22* mutant, and *unc-22* (a co-CRISPR marker for *daf-22* CRIPSR knockout) *S. hermaphroditum* were seeded onto each plate and incubated at 25C for 7 days of growth. Progeny migrate to the edge and rim of each petri dish. On days 8 and 10, nematode populations from each plate was examined and IJ was scored by morphology and viability in water. Each data point represents the percentage of IJ entry assessed from one biological replicate. Statistical analysis was performed using One-way ANOVA. The average and standard errors of six replicates from two independent experiments are shown. Numbers in parenthesis indicate total number of animals (sample size) in each condition.

## Discussion

In this study, I examined three environmental cues that influence the entry into the developmentally arrested infective juvenile (IJ) stage of the parasitic nematode *S. hermaphroditum*: symbiotic bacteria as a food source, temperature, and IJ-inducing pheromone (Table 1). These three environmental factors are well known to regulate dauer development in *Caenorhabditis* nematodes. This study shows that *S. hermaphroditum,* unlike *C. elegans,* does not require exogeneous pheromone to enter IJ stage in the lack of live symbiotic bacterial food. This discovery raises the question of whether the IJ stage represents an “alternative” developmental path, as traditionally considered, or whether it may instead represent the default developmental trajectory, with reproductive development serving as the alternative state.

**Table 1.** Environmental impact on *C. elegans* dauer and *S. hermaphroditum* IJ entry.

| Species | Bacterial food |  | Heat | Pheromone |  |
| --- | --- | --- | --- | --- | --- |
|  | Live symbiont | Killed bacteria |  | Liquid culture | mimic natural environment |
| *C. elegans* dauer | NA* | inhibit | promote | promote | NA |
| *S. hermaphroditum* IJ | inhibit | promote | inhibit | minor effect | promote |

\*NA = not available

### More than food: bacterial diet as a signal for diapause

In *C. elegans*, the presence of bacterial food, particularly heat-killed *E. coli*, acts as a competitive signal to inhibit dauer entry (Golden and Riddle 1984). *E. coli* OP50 is widely used as a food source for *C. elegans* in laboratory settings, not because it represents a natural member of the nematode’s microbiome, but because it is another model organism that is readily available and easy to culture (Frézal and Félix 2015).

Despite the similarities between the *C. elegans* dauer stage and the *S. hermaphroditum* IJ stage, the diapause programs of these two species reflect fundamentally different lifestyles outside of the Petri dish. In the wild, *C. elegans* is a free-living soil nematode that follows a boom-and-bust life cycle, experiencing unpredictable fluctuations in food sources of varying species of microbes from decaying plant stems and fruits (Fig. 1C). The distribution of food is also random in both space and time (Frézal and Félix 2015). By contrast, *S. hermaphroditum* follows a more predictable life cycle: infecting insect hosts, reproducing primarily by feeding on monoxenic culture of symbiotic bacteria, and then transmitting as IJs through the soil. Given this relatively predictable symbiotic lifestyle, *S. hermaphroditum* is likely to have evolved the ability to sense and respond to symbiont signals that forecast upcoming environmental changes (Fig. 1A). Consistent with this hypothesis, *S. hermaphroditum* exhibits a clear ON/OFF switch in its IJ developmental decision based on the viability of its symbiotic bacteria as a food source. This strategy may allow the nematode to identify the nutrient depletion stage within the insect cadaver, signaling the onset of the transmission phase. Another possibility is that *S. hermaphroditum* specifically detects signals derived from its symbiotic bacterium based on a co-evolved binary symbiotic relationship (Fig. 1A), whereas *C. elegans* may respond to more general cues from diverse microbial “food source” based on its multi-species microbiome consortia (Fig. 1B), therefore would require nematode species-specific exogeneous pheromone as the major cue for dauer formation (Figs. 2D and 2E).

Future research should investigate how microbial diets and their metabolites regulate diapause formation across different nematode species. For example, in *S. hermaphroditum*, an important question is which metabolites produced by live versus dead symbiotic bacteria promote reproduction and IJ formation, respectively (Fig. 5, section 1).

**Figure 5:**
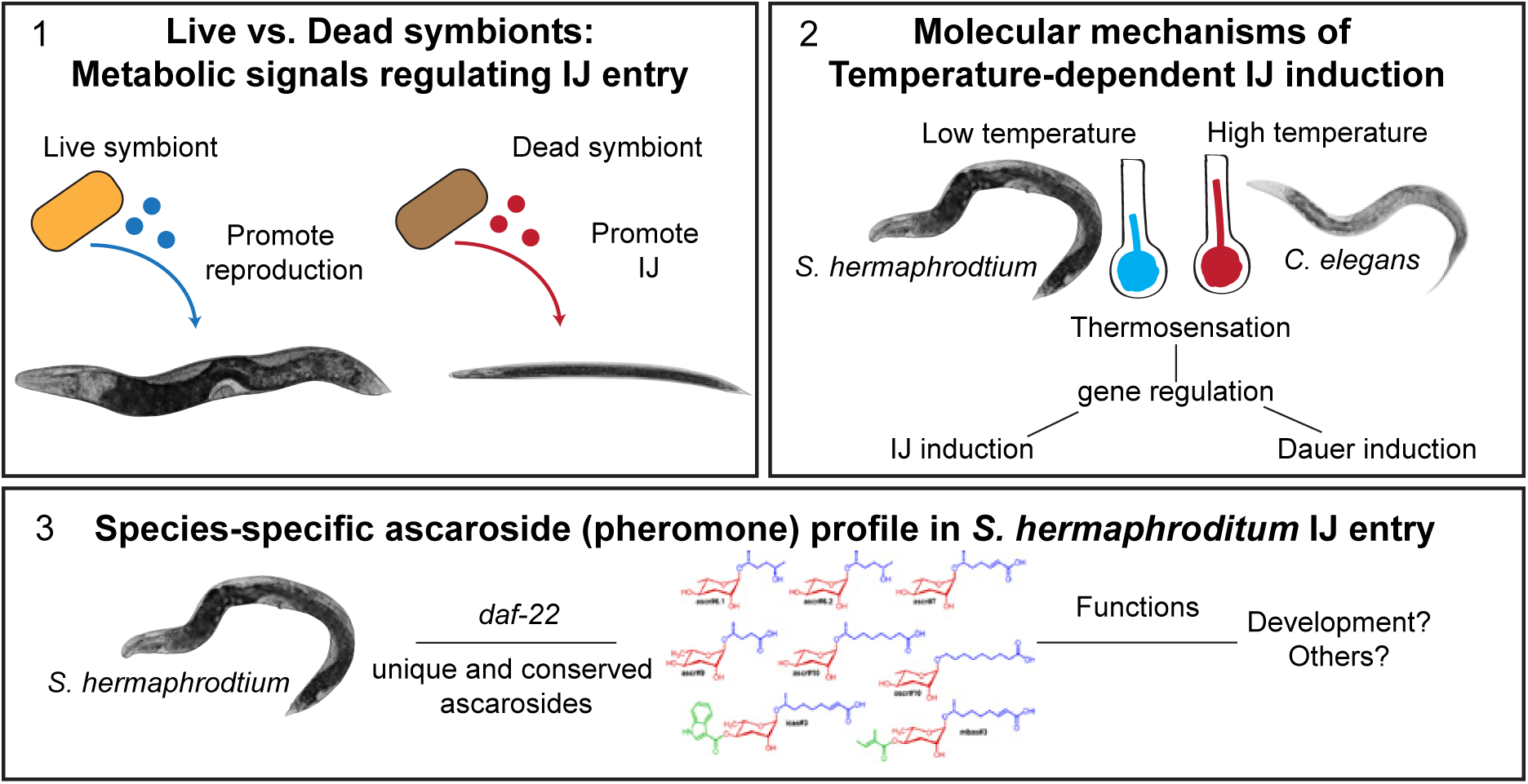
Future directions in comparative diapause development using *S. hermaphroditum* and *C. elegans* as nematode models.

### Temperature as an ecological cue: divergent thermal regulation of nematode diapause

In the paradigm of *C. elegans* dauer development, the percentage of dauer larvae formed increases at higher temperatures (Golden and Riddle 1984). One striking finding from this research is that *S. hermaphroditum* IJ formation is instead **reduced** at elevated temperatures. This effect is synergistic with pheromone presence at lower temperature. To this date, *S. hermaphroditum* strains have been exclusively isolated from tropical regions in Southeast Asia. The current strain used in this study (PS9179) was originally isolated from India and has been inbred in the laboratory for approximately ten generations (Bhat et al. 2019; Cao et al. 2021). In contrast, *C. elegans* is found worldwide, predominantly in humid temperate regions. The *C. elegans* wild-type (N2) strain used here originated from Bristol, England, a region with a mild oceanic climate, and has been cultured in laboratories for many generations before being cryopreserved (Frézal and Félix 2015). These distinct climatic origins may contribute to the species-specific differences in how thermosensory cues are perceived. A deeper understanding of these processes will require investigating the neurogenetic basis of *S. hermaphroditum* IJ development (Fig. 5, Section 2).

### Beyond *C. elegans:* decoding pheromone signals in parasitic nematodes

EPN pheromones are known to influence IJ dispersal, infectivity, and recovery (Kaplan et al. 2012, 2020; Noguez et al. 2012), but their role in dauer or IJ entry has been studied almost exclusively in *C. elegans*. In this study, I demonstrate that the *C. elegans* dauer entry assay can be adapted to investigate EPN IJ entry with modifications. One technical hurdle is that egg-laying in *S. hermaphroditum* gravid adults is significantly slower (5–6 hours to lay 50–100 eggs per 10 adults) than in *C. elegans* (1–2 hours for the same number of eggs). A second hurdle is the low potency of crude pheromone extracted from liquid culture. Various liquid culture methods for EPNs have been developed using both flasks and bioreactors (Ehlers 2001; Cortés-Martínez and Chavarría-Hernández 2020). EPN liquid cultures typically require complex ingredients, such as raw rabbit liver, agave juice, and dried egg yolk to support high-density nematode growth (20–300 IJs per μL). However, these components can be difficult to obtain. Currently, the most efficient method for EPN pheromones have been extracted from infected and spent insect cadavers. The insect cadaver-based methods may yield sufficient pheromone, with an estimated 100–200 μL of biologically relevant pheromone extractable per cadaver (Kaplan et al. 2020). In this study, I tested a simplified liquid culture system using LB medium supplemented with 5 μg/mL cholesterol and symbiotic bacteria for *S. hermaphroditum*. The nematodes fed on the symbiotic bacteria, reproduced, and reached a population density of ∼10 IJs per μL. This density is approximately 50-100% of that achieved in *C. elegans* liquid cultures used for crude pheromone extraction but still may not be sufficiently high to yield biologically relevant potency. Future work should explore simple and accessible approaches to better facilitate pheromone extraction from EPNs. Because nematode-produced pheromones function as species-specific signaling molecules that regulate physiology, future characterization of ascarosides in *S. hermaphroditum* will provide a foundation for understanding species-specific behaviors and developmental programs in these parasitic nematodes (Fig. 5, Section 3).

## Conclusion

This study establishes experimental protocols to investigate environmental factors regulating infective juvenile (IJ) formation in the parasitic nematode *S. hermaphroditum* and compares this developmental transition with dauer formation in *C. elegans*. Our data demonstrate that, unlike *C. elegans* dauer entry, which requires exogenous pheromone under laboratory conditions, *S. hermaphroditum* IJ formation can be regulated by signals derived from its symbiotic bacteria in the absence of exogenous pheromone. Furthermore, temperature influences IJ and dauer formation in opposite directions between these two nematode species. DAF-22-dependent pheromone production is required for robust IJ formation in nutrient-rich, insect cadaver-like media that mimic the natural habitat of *S. hermaphroditum*. Together, these findings reveal both conserved and divergent strategies by which parasitic and free-living nematodes perceive environmental cues to regulate developmental decisions, highlighting *S. hermaphroditum* as a promising neurogenetic model for studying parasitic nematode development.

## Acknowledgements

I thank Paul W. Sternberg and Jared Leadbetter for providing laboratory space for part of this project, Fatma Kaplan for providing valuable suggestions and comments on *C. elegans* and EPN pheromone extraction. This research was supported by Carnegie Science Endowment (MC).

## Supplemental Materials

**Figure S1:**
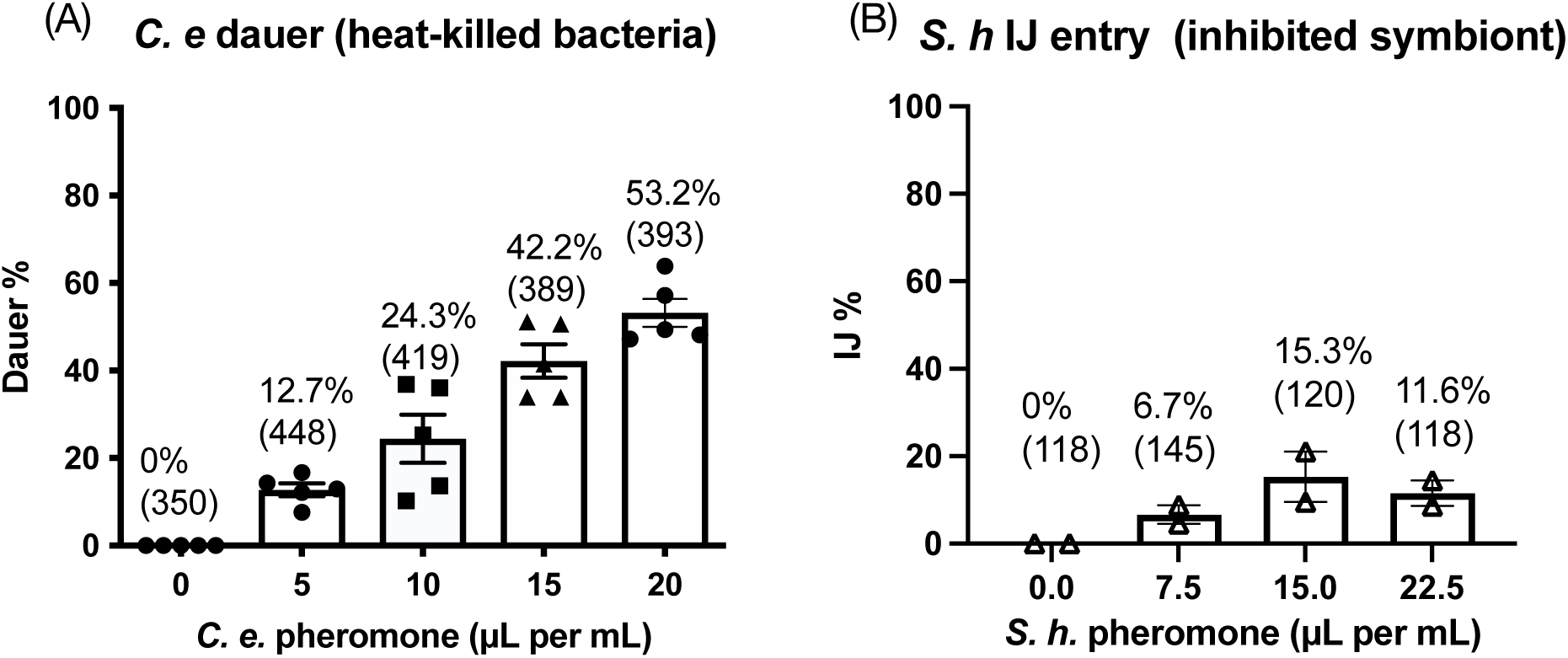
*C. elegans* dauer entry and *S. hermaphroditum* IJ entry assays on NG plates at 25.5°C. (A): *C. elegans* pheromone extracted from liquid culture induces *C. elegans* dauer formation *in vitro* in a dose-responsive manner. Each *C. elegans* pheromone plate contains 20 μL of heat-killed *E. coli* OP50 (8% w/v). (B): *S. hermaphroditum* pheromone extracted from liquid culture does not induce IJ formation in a dose-responsive manner. Each *S. hermaphroditum* pheromone plate contains 15 μL of 4% live *X. griffiniae* treated with 30 μg/mL Chloramphenicol to inhibit bacterial growth. Each data point represents the percentage of IJ entry assessed from one biological replicate. The average of percentage of dauer (A) and IJ (B) and standard errors are showed on top of each bar. Numbers in parenthesis indicate total number of animals (sample size) in each condition.

**Figure S2:**
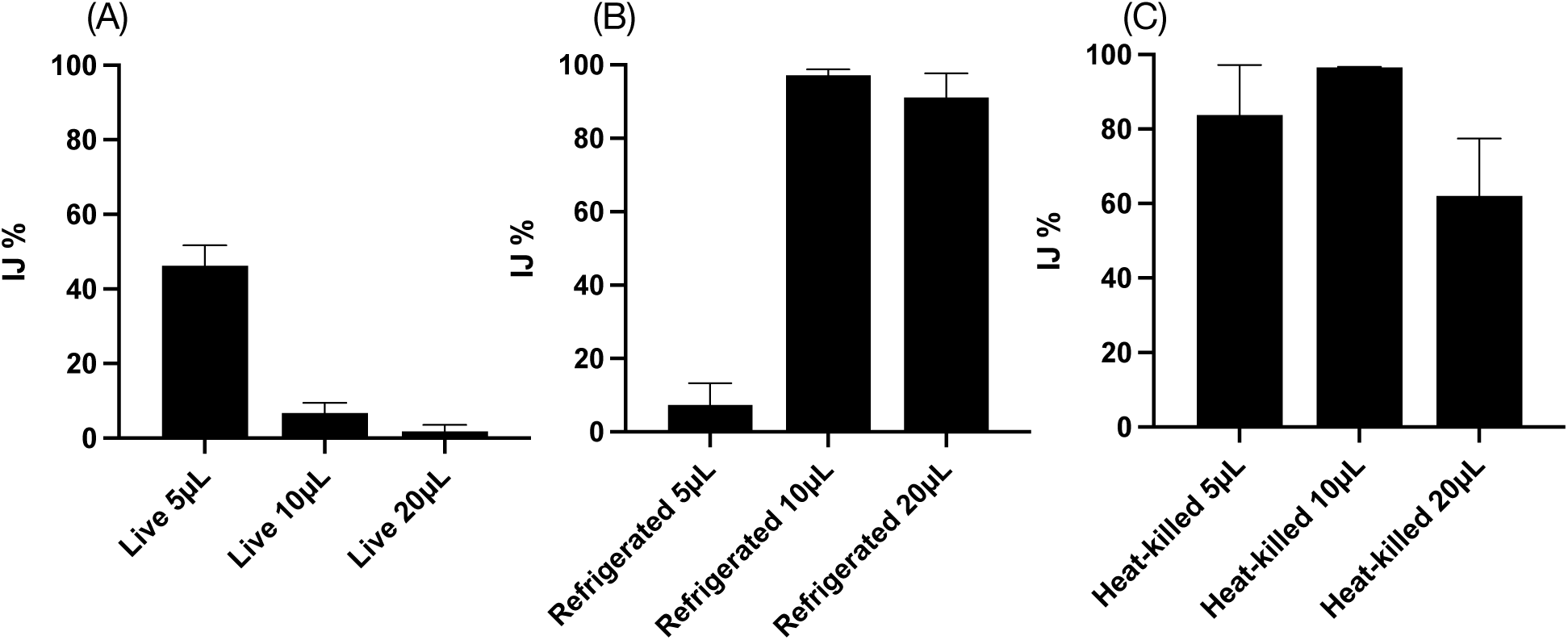
Effects of live (A), refrigerated (B), and heat-killed (C) symbiotic bacteria *X. griffiniae* on *S. hermaphroditum* IJ entry. Approximately 50-100 *S. hermaphroditum* eggs were freshly laid for each plate incubated at the 25.5°C until the IJ decision was made (24-48 hours). IJ entry on NG agar supplemented with 30 μg/mL Chloramphenicol and 5 μL, 10 μL, or 15 μL of live, refrigerated, or heat-killed symbiotic bacteria *X. griffiniae* (8% w/v). The average percent IJ entry from two biological replicates and the standard errors are shown.

**Figure S3:**
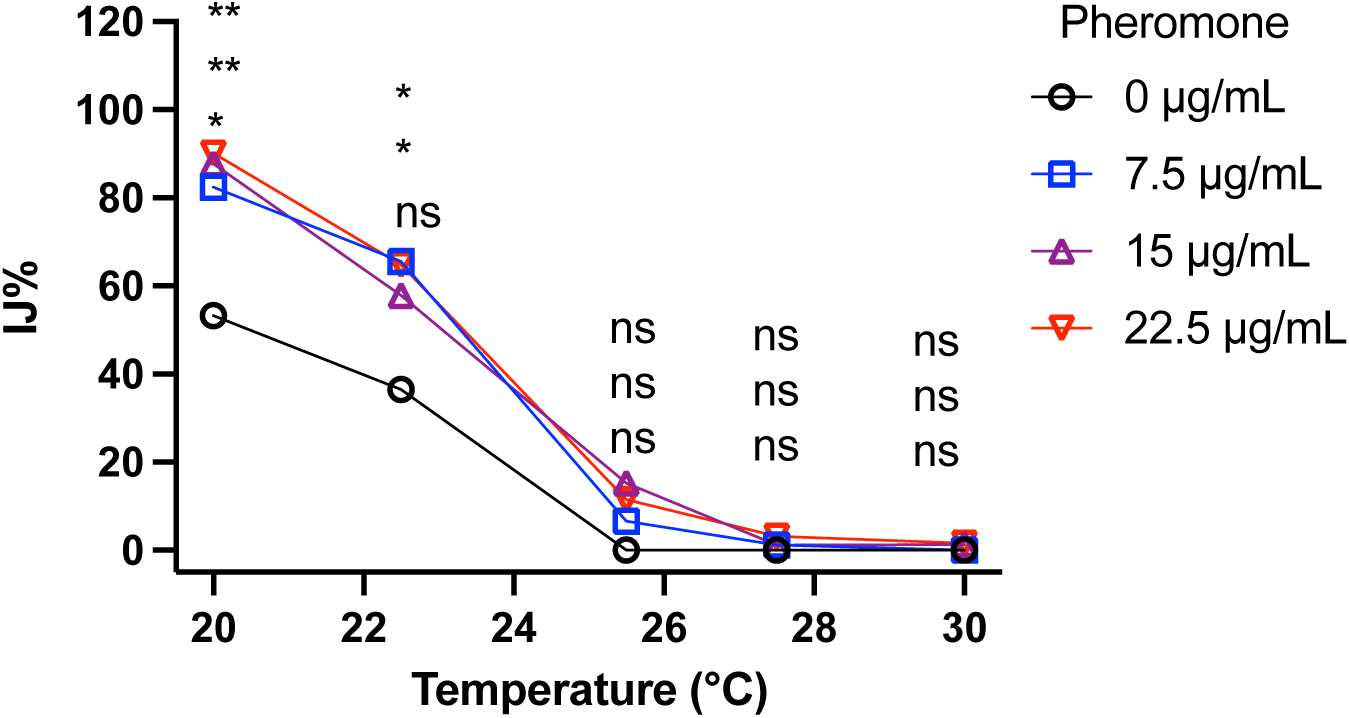
The impact of temperature on *S. hermaphroditum* IJ entry in the absence and presence of pheromone. The IJ entry assay was performed with supplementation of crude pheromone extracted from wild-type *S. hermaphroditum* liquid cultures (0 μL, 7.5 μL, 15 μL or 22.5 μL crude pheromone per mL agar). Each pheromone plate contains 15 μL of live symbiotic bacteria *X. griffiniae (*4% w/v) was freshly prepared and treated with 30 μg/mL Chloramphenicol, then added to 50-100 embryos on each agar plate. The assays were incubated at the respective temperatures (20-30°C) until the IJ decision was made in 2-4 days. Each data point represents the average of two biological replicates from the one representative independent experiment. Statistical analysis was performed using two-way ANOVA. At each temperature, data from each exogenous pheromone treatment were compared with the no-pheromone control. Statistical significance at each temperature was determined in the order that the data points appear, from the highest to the lowest percentage of infective juveniles (IJ%).

